# Time to stop; agent-based modelling of chemoaffinity with competition

**DOI:** 10.1101/2021.09.02.458703

**Authors:** Sebastian S. James, Stuart P. Wilson

**Affiliations:** Adaptive Behaviour Research Group, Department of Psychology, The University of Sheffield, Sheffield, UK

## Abstract

In the classic Chemoaffinity theory, the retinotectal axon projection is thought to use pairs of orthogonal signalling gradients in the retina to specify the eventual location of synapses made on the surface of the tectum/superior colliculus. Similar orthogonal gradients in the tectum provide a coordinate system which allows the axons to match their prespecified destination with the correct location. Although the Ephrins have been shown to guide axons toward their destination, there has yet to emerge a complete account of the local interactions which halt the axonal growth cones in the correct locations to recreate the topography of the retinal cells. The model of Simpson and Goodhill (2011) provides an account of the basic topographic arrangement of cells on the tectum, as well as reproducing well known surgical and genetic manipulation experiments. However, it suffers from the absence of a local chemotactic guidance mechanism. Instead, each agent in their model is given instantaneous knowledge of the vector that would move it toward its pre specified destination. In addition to the globally supervised chemoaffinity term, Simpson and Goodhill (2011) introduced a competitive interaction for space between growth cone agents and a receptor-ligand axon-axon interaction in order to account for the full set of experimental manipulations. Here, we propose the replacement of the chemoaffinity term with a gradient following model consisting of axonal growth cone agents which carry receptor molecule expression determined by their soma’s location of origin on the retina. Growth cones move on the simulated tectum guided by two pairs of opposing, orthogonal signalling molecules representing the Ephrin ligands. We show that with only the chemoaffinity term and a receptor-ligand based axon-axon interaction term (meaning that all growth cone interactions are by receptor-ligand signalling), a full range of experimental manipulations to the retinotectal system can be reproduced. Furthermore, we show that the observation that competition is not and essential requirement for axons to find their way (Gosse et al., 2008) is also accounted for by the model, due to the opposing influences of signalling gradient pairs. Finally, we demonstrate that, assuming exponentially varying receptor expression in the retina, ligand expression should either be exponential if the receptor-ligand signal induces repulsion (i.e. gradient descent) or logarithmic if the signal induces attraction (gradient ascent). Thus, we find that a model analogous to the one we presented in James et al. (2020) that accounts for murine barrel patterning is also a candidate mechanism for the arrangement of the more continuous retinotectal system.

## 1. Introduction

The retinotectal projection has proved to be a deep mine of information for the study of how the cells of the central nervous system are accurately connected together into functional networks. This projection connects the light-gathering cells in the retina to movement-related cells in the optic tectum (known as the superior colliculus in mammals). Light sources originating close to each other in the environment tend to activate retinal cells situated close together in the eye so that an image of the environment is formed on the retinal surface. It has been discovered that the topography of the retina is preserved within brain regions that process this information such that cells which are adjacent within the retina primarily excite cells adjacent in the tectum. This indicates that during development there must exist a mechanism which ensures the correct arrangement of the axons which leave the retina and connect to cells in the tectum.

One reason for the success of the study of the retinotectal projection is its capacity to be experimentally manipulated. In some non-mammalian species, both the retina and the tectum can be partially ablated, or even physically reorganised in-vivo, after which axons regrow to restore the order and function of the system for the individual animal. This manipulability was exploited in influential work by R. W. Sperry and co-workers during the mid-twentieth century, leading to Sperry’s 1963 summary of the *chemoaffinity theory* (Sperry, 1963) which proposes the existence of morphogenetic gradients that guide axons to their destination. The chemoaffinity theory was given robust support by the discovery of the ephrin ligands and their receptors (Cheng et al., 1995; Drescher et al., 1995) which have been shown to form into graded expression fields in the retina (Braisted et al., 1997), tectum (Braisted et al., 1997; Feldheim et al., 2000) as well as in other sensory systems, such as the somatosensory system (Vanderhaeghen et al., 2000). The Ephrin ligands have a clear effect on axonal outgrowth, as shown in in-vitro (Cheng et al., 1995; Drescher et al., 1995; Hansen et al., 2004) and in-vivo (Frisén et al., 1998; Rodger et al., 2000; Mann et al., 2002; Hindges et al., 2002) studies.

It is tempting to consider the retinotectal projection well understood. With a comprehensive theory supported by a biochemical mechanism, is there anything left to understand? That question can be answered by reviewing retinotectal modelling papers.

Mini-review here, which contrasts some of the modelling papers. The upshot is that a central problem with models is not how the axons know how to get closer to their destination, but how they know they have *arrived* at their true destination. Also make the point that many of the models are phenomenological in nature and do not elucidate the mechanisms behind the arrangement.

In a recent paper, we proposed a self-organising mechanism, based on morphogenetic signalling gradients, which can arrange axons growing from the thalamus to the somatosensory cortex into the well known murine barrel cortex pattern (James et al., 2020). In characterising that system, we explored the effect of various types of noise. We found that the mechanism was robust to noise in the expression of the signalling molecules over a wide range of amplitudes and length-scales, and that noise in the interaction parameters, which are obtained by sampling the signalling molecules in the source tissue (the thalamic barreloid field) could cause topological defects. The question of noise in axon guidance has been explored by Goodhill (2016).

### New introduction

Simpson and Goodhill presented a model in which the chemoaffinity mechanism is non-biological. We offer a model in which a competition mechanism counteracts the drive of axons along the ephrin gradient. Will we need the axon-axon interaction mechanism based on non-same-Eph repulsion?

### Another approach

We present a model of gradient-following based on graded interactions between the growth cones of retinal ganglion cell axons and the tectal surface. We show that a combination of exponentially graded expression in the retina (Reber et al., 2004) and exponentially (for repulsively interacting receptor ligand pairs) *or* logarithmically (for attractively interacting receptor/ligands) graded expression on the tectum permits a purely local model to reproduce the majority of reported phenomena associated with retinotectal organisation.

There have been many approaches to modelling the organisation of the retinotectal projection. Many of these models are wholly or partially phenomenological.

A model which carefully deals with receptor binding is given by Naoki (2017) (see also Mortimer et al., 2009).

Gradient based models: Nakamoto et al. (1996)

## 2. Model

Our model was inspired by the agent-based model of Simpson and Goodhill (2011) and, in common with that work, consists of agents representing axonal growth cones, originating from a square retina and moving upon a square tectum of side length 1 in arbitrary units. Simpson and Goodhill (2011) model three contributions to the movement of the growth cones; chemoaffinity, competition for space and axon-axon interactions. The first two of these are modelled without reference to a particular mechanism; each agent is provided with globally-supervised information about its instantaneous location and its target location, allowing it to move along a vector towards the target. Competition is implemented as a movement of constant distance along the vector between two agents if those two agents are within a threshold distance of one another. Interaction is signal-induced repulsion, which is triggered if two agents are both within the threshold distance of each other and have compatible EphA receptor expression.

In our model, space-based competition was retained (though sometimes omitted) and chemoaffinity and axon-axon interactions were remodelled so that all interactions occurred via signal transmission between receptors and ligands, both of which are expressed in graded patterns on both retina and tectum. Signals induce attractive or repulsive interactions; for chemoaffinity the interaction is to ascend or descend the gradient of ligand expression on the tectum; for axon-axon interactions, the strength of short-range repulsion (or attraction) is governed by the receptor-ligand expression on pairs of axons.

### Receptor and ligand expression

Each retinal ganglion cell projects *n* growth cone agents (referred to as *branches*) which carry a set of 4 receptors, *r*_*i*_, and 4 ligands, *l*_*i*_, indexed by *i* and expressed at levels determined by the cell’s location on the retina. The expression levels of the receptors and ligands vary with respect to the cell’s position on the retinal surface. Each receptor or ligand varies with respect only to one dimension. Receptor expression gradients are arranged in orthogonal pairs with the gradient of *r*_0_ being orthogonal to that of *r*_1_. *r*_2_, whose gradient is opposite to *r*_0_, is orthogonal to *r*_3_. Ligands are also arranged in orthogonal pairs, with *l*_0_ opposing *r*_0_ (as *r*_0_ increases, *l*_0_ decreases) and *l*_1_ orthogonal to *l*_0_ and opposing *r*_1_.

Because there is convincing evidence that EphA and EphB receptors are expressed in exponentially increasing patterns (Reber et al., 2004; Feldheim et al., 2000; Brown et al., 2000; Koulakov and Tsigankov, 2004), we use an exponential form for retinal receptor expressions, in common with other modelling studies (Reber et al., 2004; Koulakov and Tsigankov, 2004; Simpson and Goodhill, 2011). We adopt the same precise form for the retinal receptor expression as Simpson and Goodhill (2011) which is

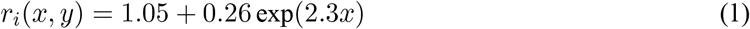

The simulated tectum expresses the same types of receptors and ligands, also in orthogonal pairs of gradients. In this work, we did not consider the effect of reverse signalling from tectal receptors to RGC ligands so we only modelled tectal ligand expression, *L*_*i*_. Although several studies model tectal ligand expression with exponential functions (Koulakov and Tsigankov, 2004), the experimental evidence for ligand expression is more ambiguous (Supp. Fig **X**). Correspondingly, we leave open the possibility that tectal ligand expression may be modelled by exponential, linear or logarithmic functions:

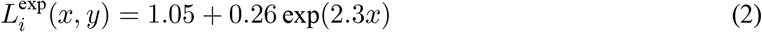

or

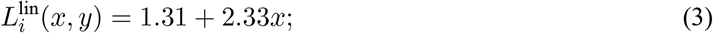

or

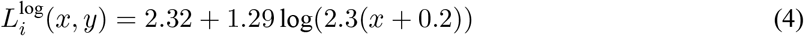

Each function has approximately the same expression values at *x* = 0 and *x* = 1. Opposing functions are obtained by substituting (1 *− x*) for *x*.

While we do not explicitly name *r*_0_, *r*_1_, etc. as *EphA, EphB*, the suggestion is that *r* includes EphA, EphB, Ryk (Schmitt et al., 2006) and Neogenin (Rajagopalan et al., 2004) receptors and that *l* and *L* include the ephrin-A, ephrin-B, Wnt3 (Schmitt et al., 2006) and RGM (Monnier et al., 2002) ligands, each of which has been shown to play a role in retinotectal map formation.

Figure 1 shows colour maps of receptor and ligand expression, in this case tectal ligand expression maps are linear. Note that the expression is discretized, and in simulations, the gradient is computed numerically from the discretized expression. This makes it straightforward to perform tectal and retinal ‘graft’ manipulations. Coordinates within the simulated retina and tectum are normalised, with 1 corresponding to 2 mm on the tectum and on the retina for mouse (Reber et al., 2004). This means that growth cones of approximate radius 5 *µ*m = 0.005 mm (**?**) have a radius *r* = 0.0025 in the arbitrary units of the simulation. For a realistic simulation based on mouse, the approximate RGC density in the retina is around 44000 cells per retina (**?**). This sets the side length of the retina to 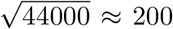. This results in a simulation of prodigious computational size, which although tractable (10 s per step, requiring several hours, rather than days to run) is not convenient. To maintain the relationship between growth cone size and cell density, we permit a smaller number of growth cones (400 *×* 8 = 3200) but keep the fractional area of the branches at 0.86 (44000 *× πr*2) (if there is one cone per axon) or 6.9 (if there are 8 cones per axon).

**Figure 1:**
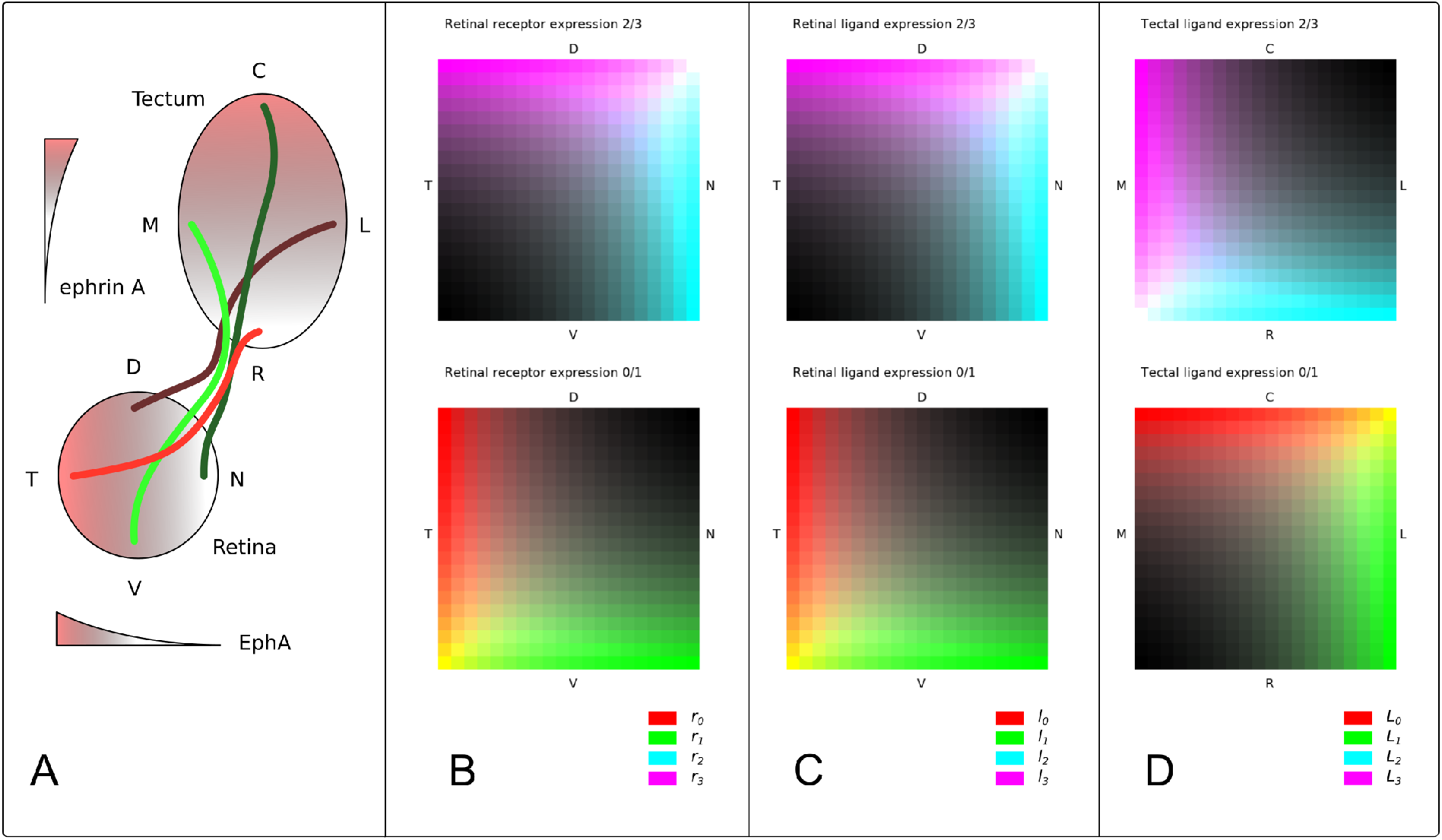
Simulated retinal and tectal signalling molecule expression. Four receptor types are expressed, and four corresponding ligand types. Receptors of type *i* are activated by ligands of type *i*. **A** Retinal receptor expression, *r*_*i*_. Four receptors are expressed and displayed here in two dual-colour maps. *r*_0_ and *r*_1_ are red and blue; *r*_2_ and *r*_3_ are cyan and magenta. *r*_0_ increases as an exponential function of *−x*; *r*_1_ increases with an exponential function of −*y. r*_2_ and *r*_3_ have the opposite sense. Retinal receptor expression is modelled with exponential functions because there is ample evidence to support this form. *N*: nasal, *T*: temporal, *D*: dorsal, *V*: ventral. **B** Retinal ligand expression, *l*_*i*_, is expressed in complimentary gradients (Hornberger et al., 1999). **C** Tectal ligand expression *L*_*i*_. Tectal ligand expression is shown in a linear form, as there is less evidence to determine whether ligand expression is exponential, linear or logarithmic on the tectum. *L*: lateral, *M*: medial, *C*: caudal, *R*: rostral.

### Movement

At each timestep in a simulation, the position, **x**_*b,t*_, at time *t* of each branch *b* is updated according to

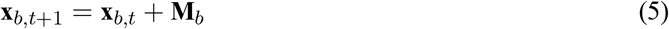

where **M**_*b*_, the movement vector, is the sum of a chemoaffinity effect, **G**_*b*_, axon-axon competition, **C**_*b*_, an axon-axon receptor-ligand interaction, **I**_*b*_ and a ‘border effect’, **B**_*b*_, which acts to retain branches within the tectal region:

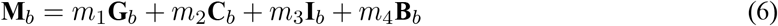

evaluated at time *t. m*_1_–*m*_4_ are scalar parameters.

### Chemoaffinity

In this model, the signal transmitted when a ligand binds to a receptor on a branch (forward signalling) can lead to one of two effects; the signal may induce an attraction towards the ligand-expressing region or a repulsion away from it. For axon-tectum interactions, we implemented attraction as a climbing of the tectal ligand expression gradient and repulsion as gradient descent. To determine the strength of the effect we assumed a purely linear receptor binding model, and set the chemotactic movement vector of the branch *b* at location **x**_*b*_ on the tectum to be

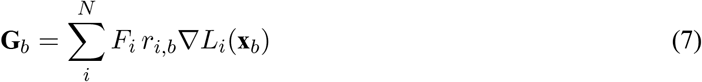

where *r*_*i,b*_ is the receptor expression on branch *b* for ligand-receptor pair *i*, ∇*L*_*i*_ is the gradient of expression of ligand *i* on the tectum and *F*_*i*_ denotes the direction of the interaction induced when a molecule of ligand *I* binds to a receptor *i* molecule. *F*_*i*_ takes the value −1 for a repulsive interaction or 1 for an attractive interaction.

### Axon-axon competition

Competition, in the context of this model, is a repulsive effect which acts between any two branches, whether they originate from different retinal ganglion cells or from the same cell. It assumes that there is some kind of self-other signal (**be sure to discuss possible origins**) which acts to discourage two cells from approaching each other.

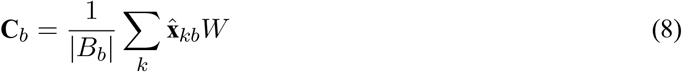

where the distance based weight *W* is given by

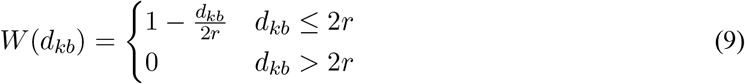

for two growth cones of radius *r* separated by a distance *d*_*kb*_. *B*_*b*_ is the set of growth cones within a distance 2*r* of branch *b*.

### Axon-axon interactions

For axon-axon interactions, repulsion causes a movement of branch *b* along a unit vector, 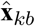, from branch *k* to branch *b* causing them to move further apart; attraction causes the opposite movement. The interaction, **I**_*b*_, acting on branch *b*, is given by

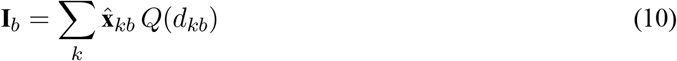

where *Q*(*d*_*kb*_) is the signalling strength between two growth cones of radius *r* a distance *dkb* from one another, given by

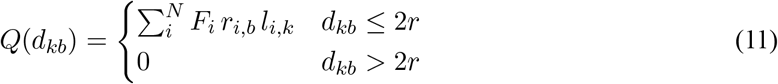

The sign of *Q* (which is dependent on the values of *F*_*i*_) determines whether the interaction, **I**_*b*_, is repulsive or attractive. *l*_*i,k*_ is the expression of ligand *i* on branch *k*.

### Border effect

We implement a border effect based on gradient following by assuming that there is some other molecular signal which acts on all branches near the boundary of the tectal tissue. For a branch with position (*x, y*), **B**_*b*_ is given by:

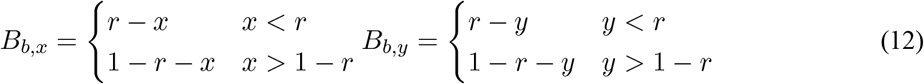

Thus **B**_*b*_ is equivalent to the action of a repulsive signalling molecule expressed around the border of the tectum, whose expression increases quadratically outside the tectum and affects any branch touching (or outside) the boundary

### Initial conditions

Branches were randomly distributed in a stripe at the rostral side of the tectum at the start of each simulation. Each RGC axon was assigned a random initial position coordinate:

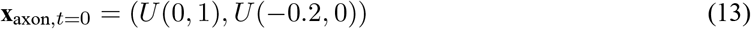

where U(p,q) is a number selected from a random uniform distribution in the range [*p, q*). Each of the *n* branches per RGC axon was given its parent axon’s initial position, plus a randomly generated offset with coordinates derived from a normal distribution of mean 0 and standard deviation 0.1.

## 3. Results

### Normal development

To enable a simulation it is necessary to make assumptions about the nature of the receptor-ligand signalled interactions (i.e. whether they are attractive or repulsive) and about the pattern of ligand expression on the tectal surface. We assume that i) All receptor-ligand signalled interactions are repulsive, as for EphA-ephrinA coupling (Drescher et al., 1995; Nakamoto et al., 1996); and ii) all tectal ligand expressions vary exponentially along a given axis, following Eq. 2. The results of running such a system are presented in Fig. 2. Fig. 2Ai shows a dual-colour map of retinal ganglion cell positions on the square retina, with red encoding position on the *x* axis (temporal to nasal) and green encoding position on the *y* axis (ventral to dorsal). Fig. 2Aii presents a fishnet plot of the expected location on the tectal surface of each RGC at the end of the development process. Thus, the ‘red’ RGC from the ventro-nasal corner should map to the caudo-medial corner of the tectum and the ‘green’ RGC from the dorso-temporal corner should find its way to the rostro-lateral corner on the tectum. The lines between the dots of Fig. 2Aii indicate the adjacency relationships between cells. Thus, colour in each panel indicates a branch or axon’s location of origin on the retina. Fig. 2B shows the initial state of the system, with branches stochastically arranged around the rostral edge of the tectum. Each dot represents the centroid of *n* = 8 branches per RGC axon. Figs. 2C and 2D show the development of the system with time, with Fig. 2D showing a well ordered arrangement, with a majority of adjacency relationships matching the experimental prediction (i.e. there are few crossed lines). The intermediate timepoint at *t* = 40 indicates that branches quickly cover the simulated tectal surface, with final adjacency relationships developing over a longer timescale. This dynamic behaviour is also evident in Fig. 2E, which shows **RMS error or some metric** of the axon centroids with respect to the experimental prediction. Fig. 2F shows the final location of each branch in the simulation, which gives a more continuous view of the order present in Fig. 2D. Finally, Fig. 2G shows the branches and their centroid path history for five selected RGCs, with path histories strongly resembling those presented by Simpson and Goodhill (2011).

**Figure 2:**
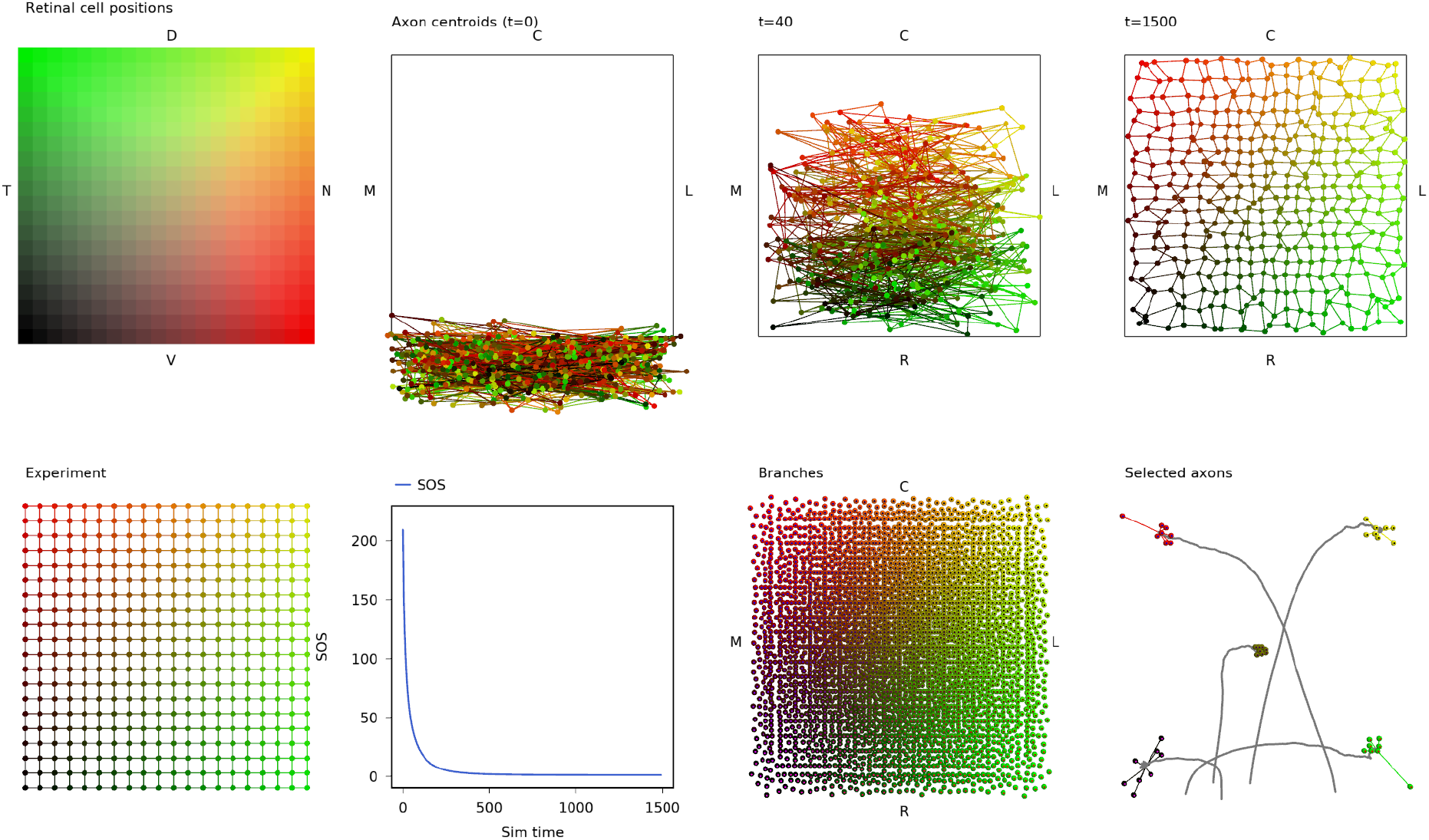
Gradient following model with axon-axon interactions.

This result shows that a gradient following model based on receptor-ligand interactions and incorporating axon-axon repulsion is able to generate the basic mapping observed in the retinotectal projection.

### Manipulations

To determine if the model can explain the many surgical and genetic manipulations described in the literature (for a review, see Goodhill and Richards, 1999) we performed simulations matching those in Simpson and Goodhill (2011) in which we either rotated or translocated grafts of the tectum; ablated portions of the retina and/or the tectum; or manipulated the receptor or ligand expressions in the model.

#### Surgical Manipulations

Fig. 3 shows the result of simulating three surgical manipulations alongside the wildtype result, which is shown on Row A. Rows B and C show graft rotation simulations (90° and 180°). In each case an 8×8 square on the tectum (each unit is 0.05×0.05) is rotated. The rotational sense can be observed clearly in the ‘Branches’ column; for the 90° rotation, the top-right (caudal-lateral) corner of the rotated square is red, whereas the top-left corner of the wildtype tectum is red. **etc**.

**Figure 3:**
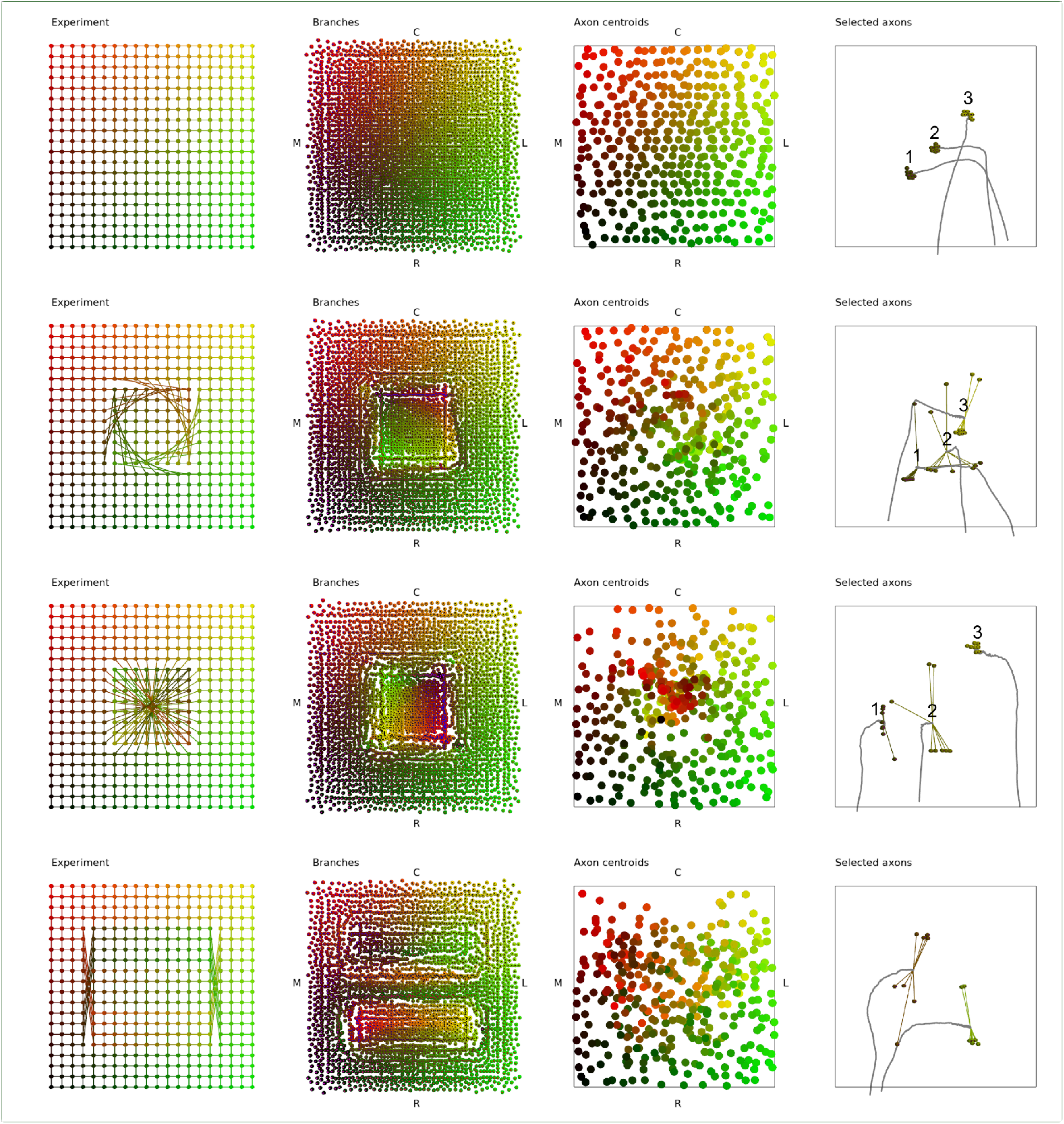
Surgical manipulations. Row A: wildtype. Row B: 90 degree rotation. Row C: 180 degree rotation. Row D: Graft swap.

Fig. 4 shows retinal ablation, tectal ablation, mismatch and compound eye experiments (Stuart: see Simpson and Goodhill Fig. 4).

**Figure 4:**
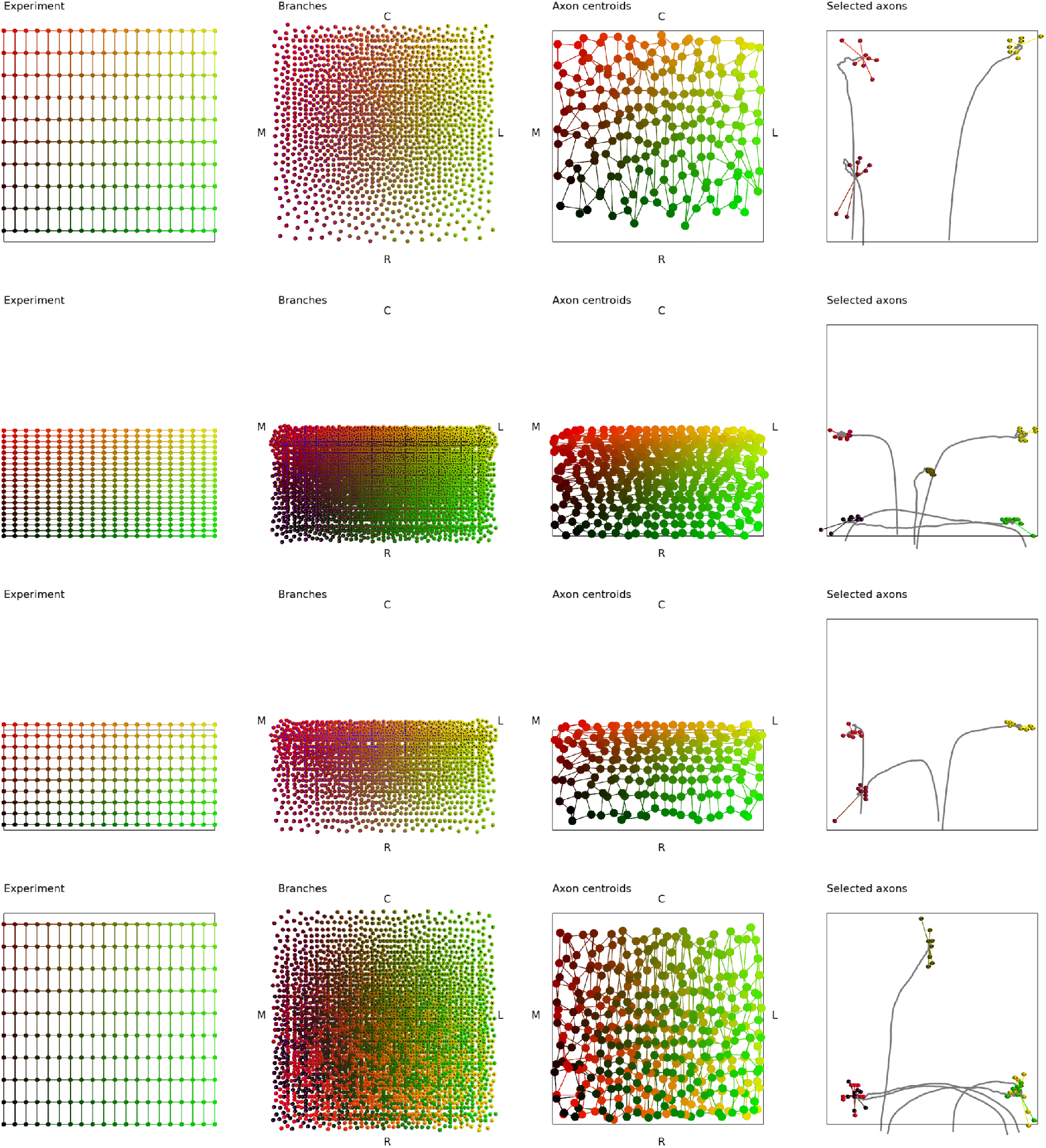
Surgical manipulations 2. Row A: Retinal ablation, Row B: Tectal ablation, Row C: Mismatch (ablate retina and also the tectum that matches the left-behind retina), Row D: Compound eye (Two nasal retinal halves ‘glued together’)

#### Genetic manipulations

Brown et al. EphA knock-in.

Describe Reber and other gene knock-in and knock-out experiments.

### ligand expression profiles

A section to show the effect of assuming linear or logarithmic expression profiles. Especially that linear expression fails to reproduce the rotation and graft swap manipulations. Hmm, actually, it looks ok?!

### Attractive receptor-ligand interactions

Show that, if you switch one of the receptor-ligand interactions in the system to be attractive instead of repulsive, then you have to make the ligand expression form logarithmic instead of exponential.

### Is chemoaffinity alone sufficient to explain the retinotectal projection?

Introductory sentence.

### FIXME: Find the expression functions for the tectum which give a decent WT result. Perhaps I already did this? Experiment with these and axon-axon with smaller radius of interaction

We now describe the behaviour of a model which has only the gradient-based chemoaffinity effect and no axon-axon interaction or border effect (i.e. *m*_2_ = *m*_3_ = 0). This model was run with *n* = 8 branches per RGC axon. All receptor-ligand signals induced repulsive interactions (i.e. gradient descent). Receptor and ligand expressions were as shown in Fig. 1, noting that tectal ligand expression was of exponential form (Eq. 2). Fig. 9A gives a colour map of retinal ganglion cell soma locations, with the colour red encoding *x* position on the temporal-nasal axis (with red signifying nasal positions). Green encodes *y* position on the dorso-ventral axis. Fig. 9B shows the expected final location of each RGC axon, according to experimental observations, in a ‘fishnet’ plot which draws lines between adjacent axons giving a visual indication of the topology of the arrangement. Fig. 9C shows the position of 400 branches at *t* = 500 (in arbitrary time units). Considering each coordinate axis separately, each branch interacts differentially with two opposing gradients. Branches which settle nearest the rostral edge of the tectum interact more strongly with the gradient *L*_0_ than with *L*_2_ (Fig 1C). A similar pair of gradients separate branches along the M-L tectal axis. The branches become positioned on a grid due to the discretization of the tectal ligand gradient (as the number of branches and tectal elements tends towards infinity, the arangement tends towards a continuous map—see Fig. S1 **FIXME**).

**Figure 5:**
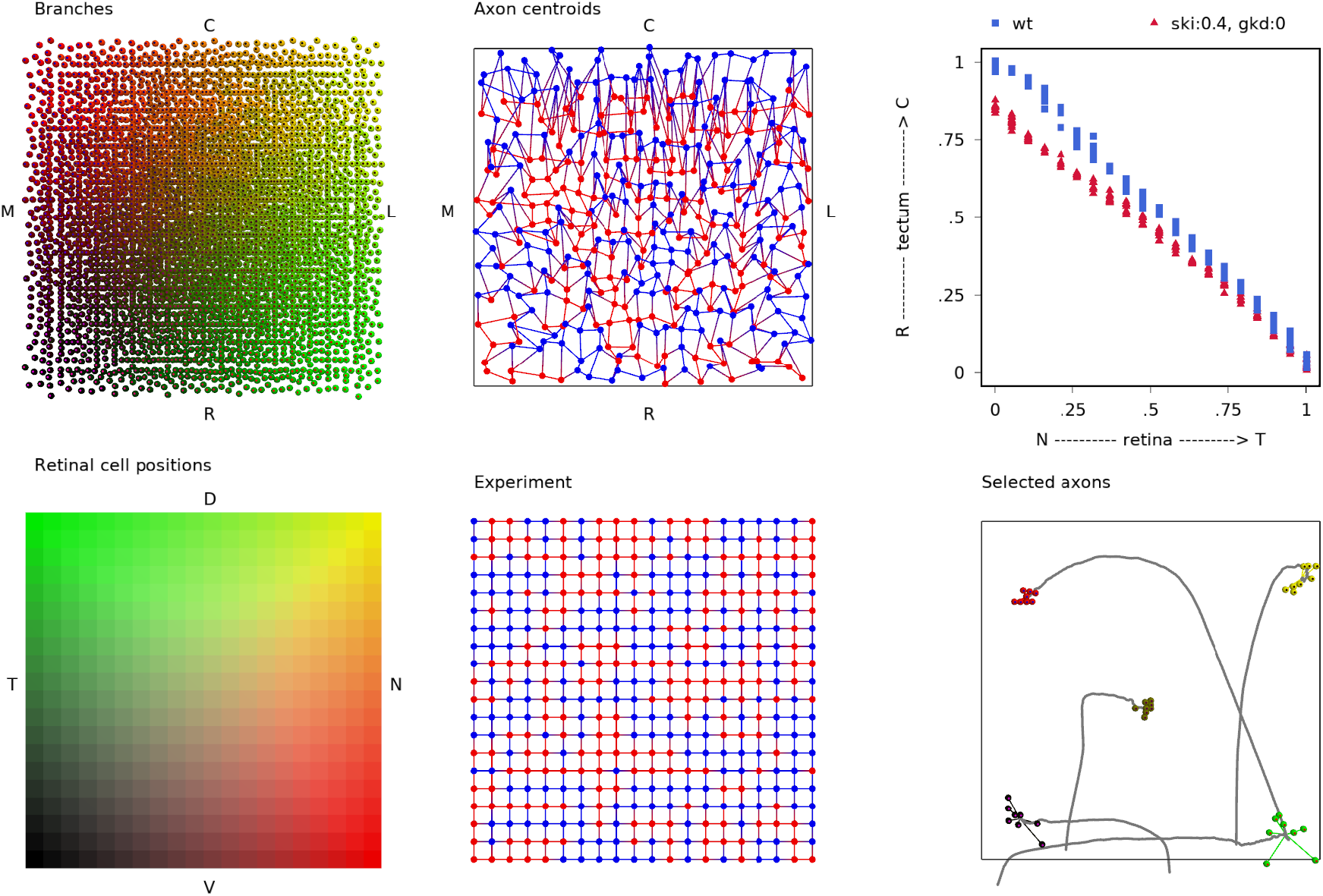
Brown et al simulation. This is EphA3 knockin - it adds a constant to half of the RGC’s *r*_0_ (randomly selected). Compare with S&G Fig. 6.

**Figure 6:**
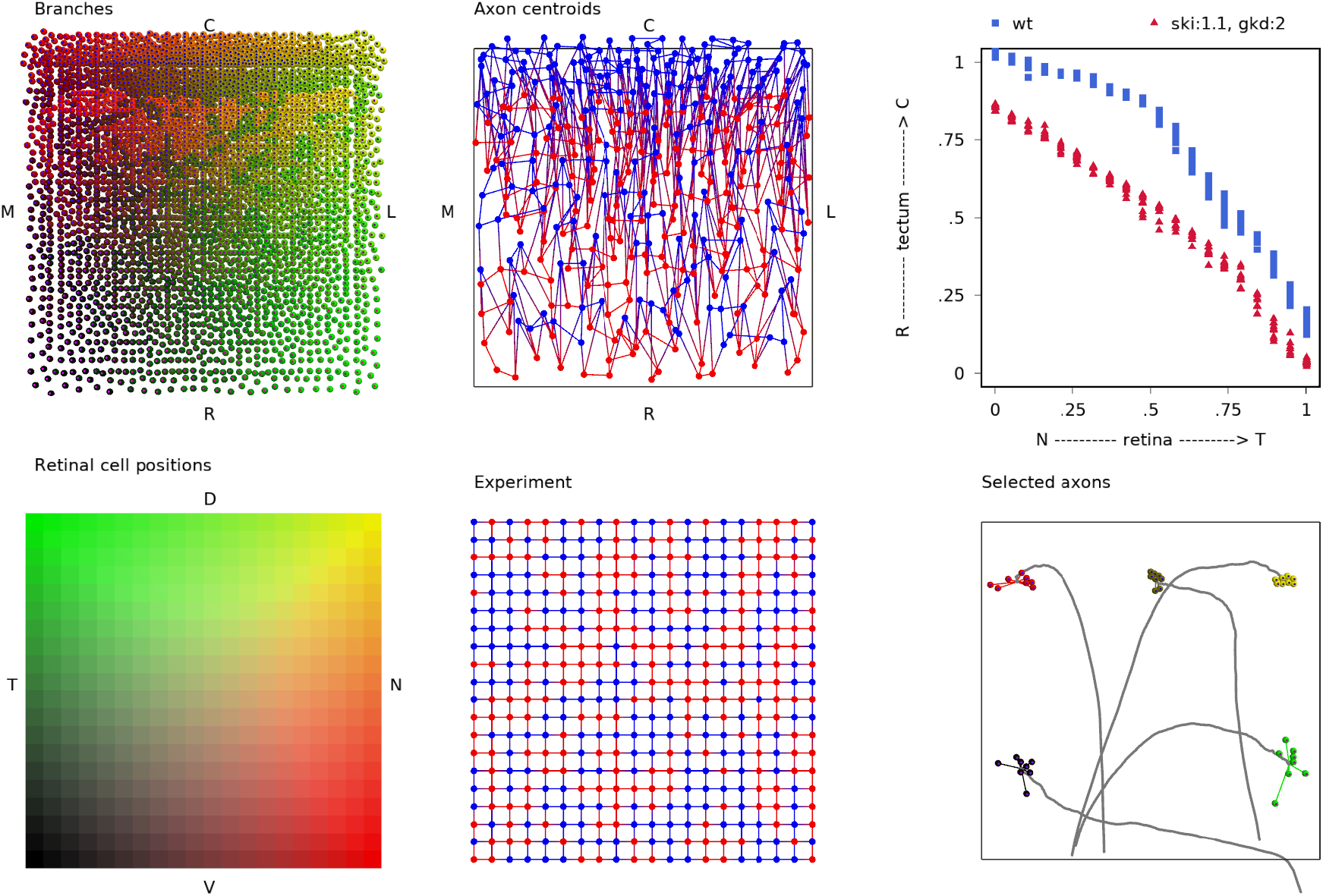
Reber et al simulation. Follow on to Brown et al (same group). Combines the selective EphA3 knockin with a uniform EphA4 knockdown.

**Figure 7:**
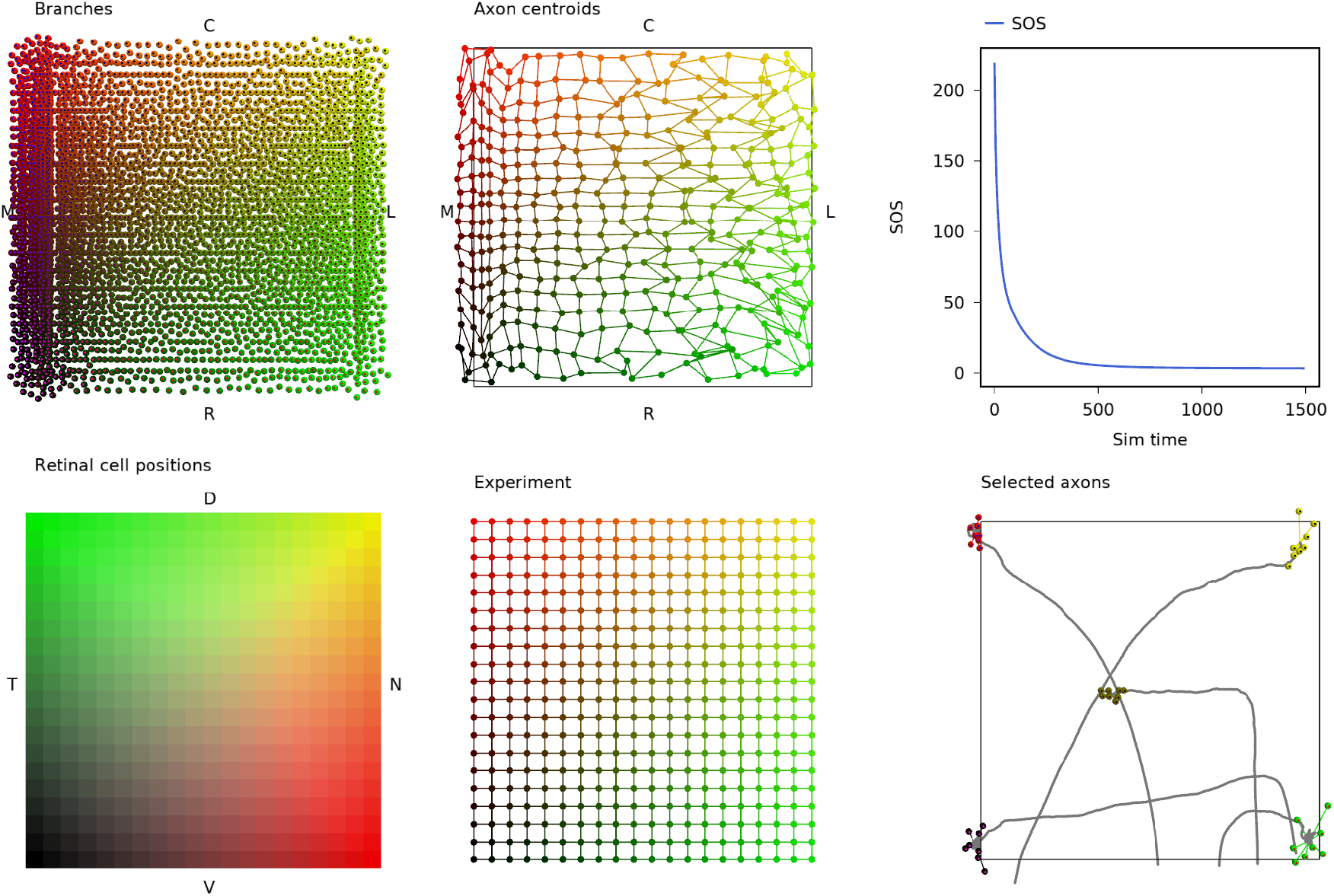
The effect of making one interaction attractive. This models switching one receptor/ligand pair to be an analogue of EphB, but retains an assumption that the ephrin-B ligand has exponential form. The change compared with the model of Fig. 2 is that the tectal ligand direction of *L*_1_ is reversed and the interaction of *r*_1_ and *L*_1_/*l*_1_ is made attractive. *L*_1_ retains the assumption of exponential change. Note the ‘bunching up’ of the field to the medial side of the tectal sheet.

**Figure 8:**
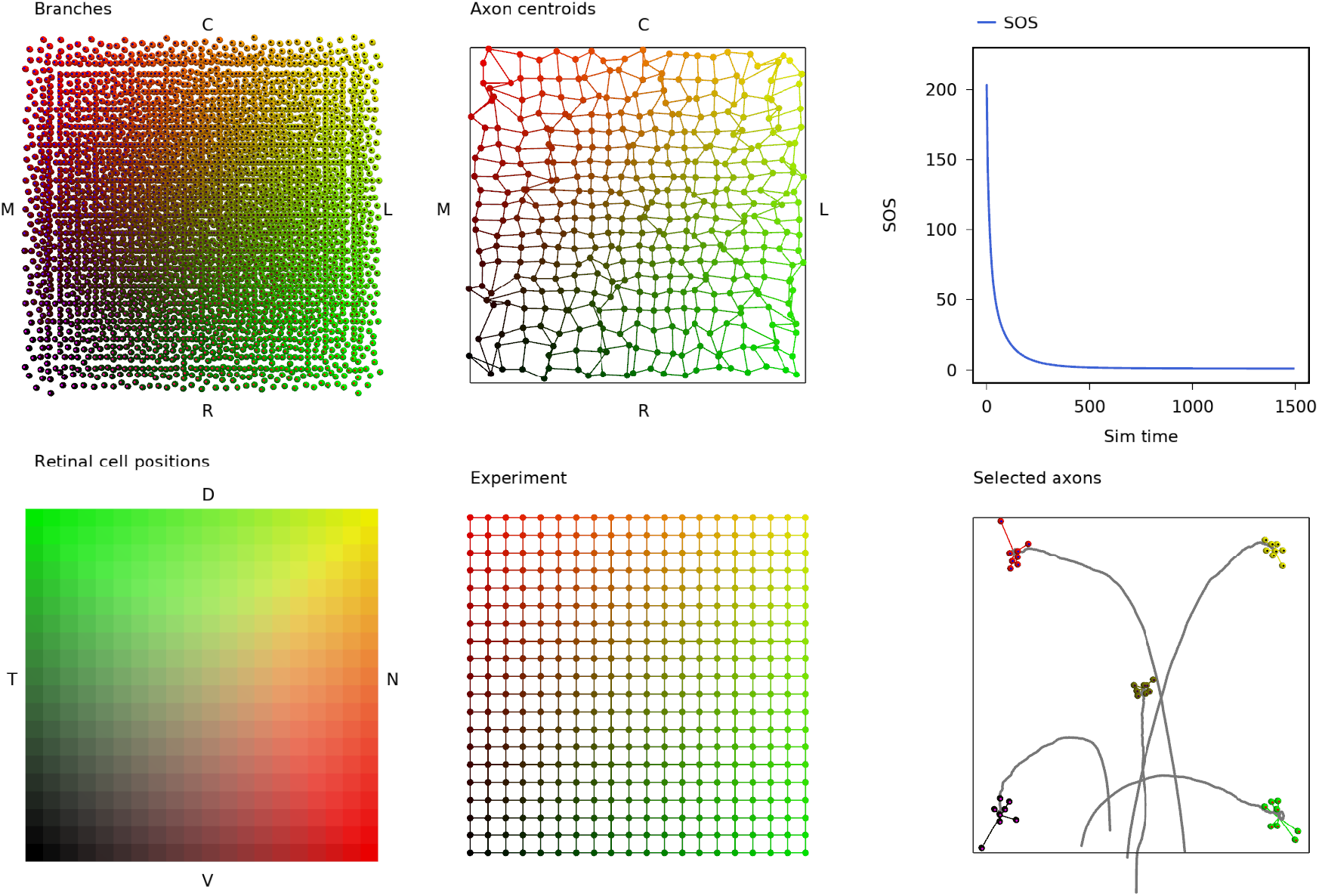
The arrangement is restored by making the expression of *L*_1_ on the tectal sheet change logarithmically.

**Figure 9:**
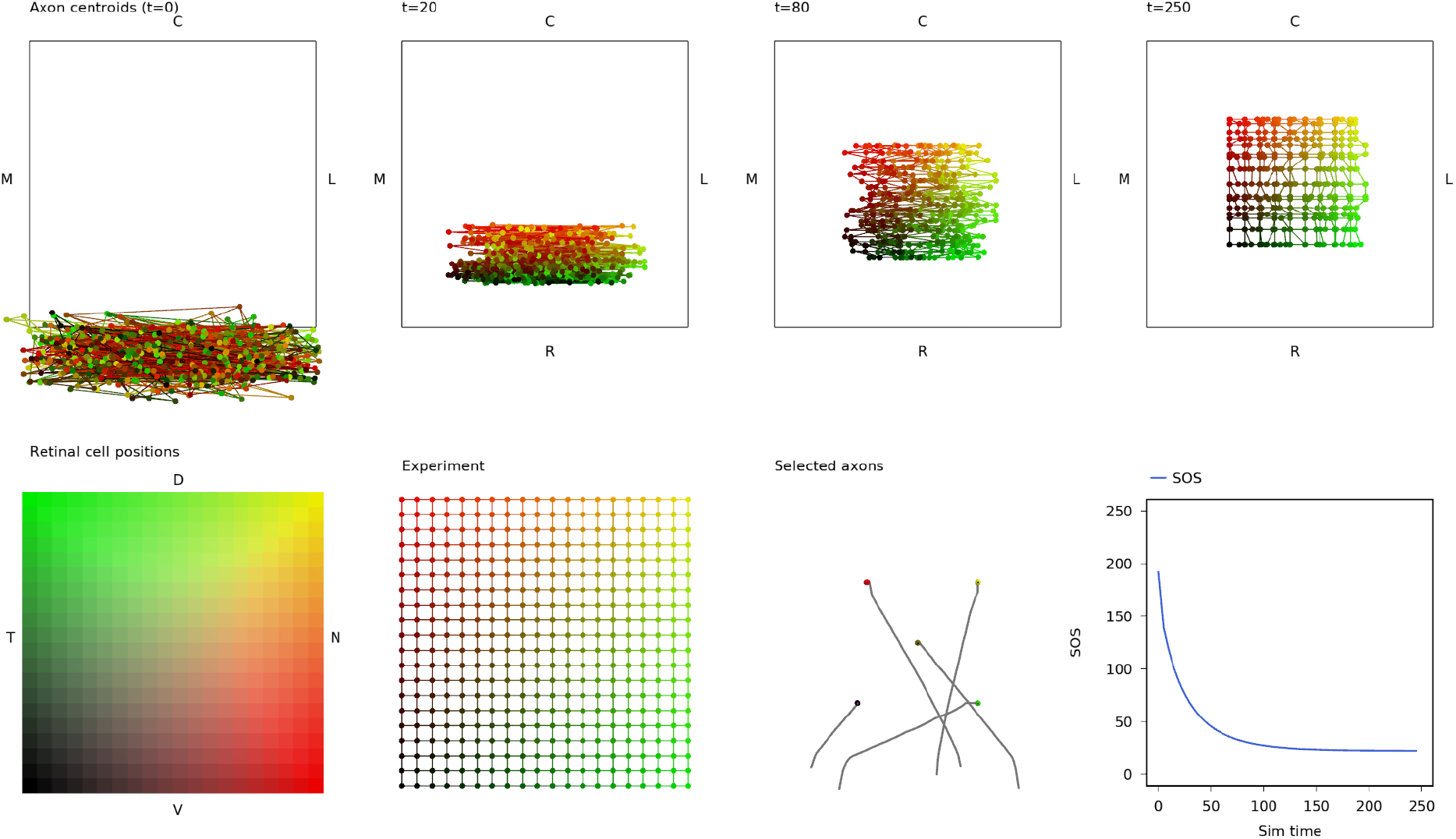
Chemoaffinity model

Fig. 9D shows a fishnet plot of the RGC axon centroids; the mean location of *n* branches per axon. Because all *n* branches for each RGC have identical receptor expression and they experience only the chemoaffinity effect, each branch attains almost the same final position and so the centroid graph strongly resembles Fig. 9B. Fig. 9D indicates that the topology of the arrangement resembles that of the experimental result, but the area over which the branches arrange themselves is limited to about 50% of the tectum. It is possible to tailor the tectal ligand expression, whcih adjusts the relative contributions of the opposing interactions, to tune the final map. This allows the chemoaffinity model to attain a good match to the experimental arrangement (Fig.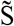2 **fixme**). However, the match is highly dependent on the form of receptor/ligand expression in both retinal and tectal cells.

**Figure 10:**
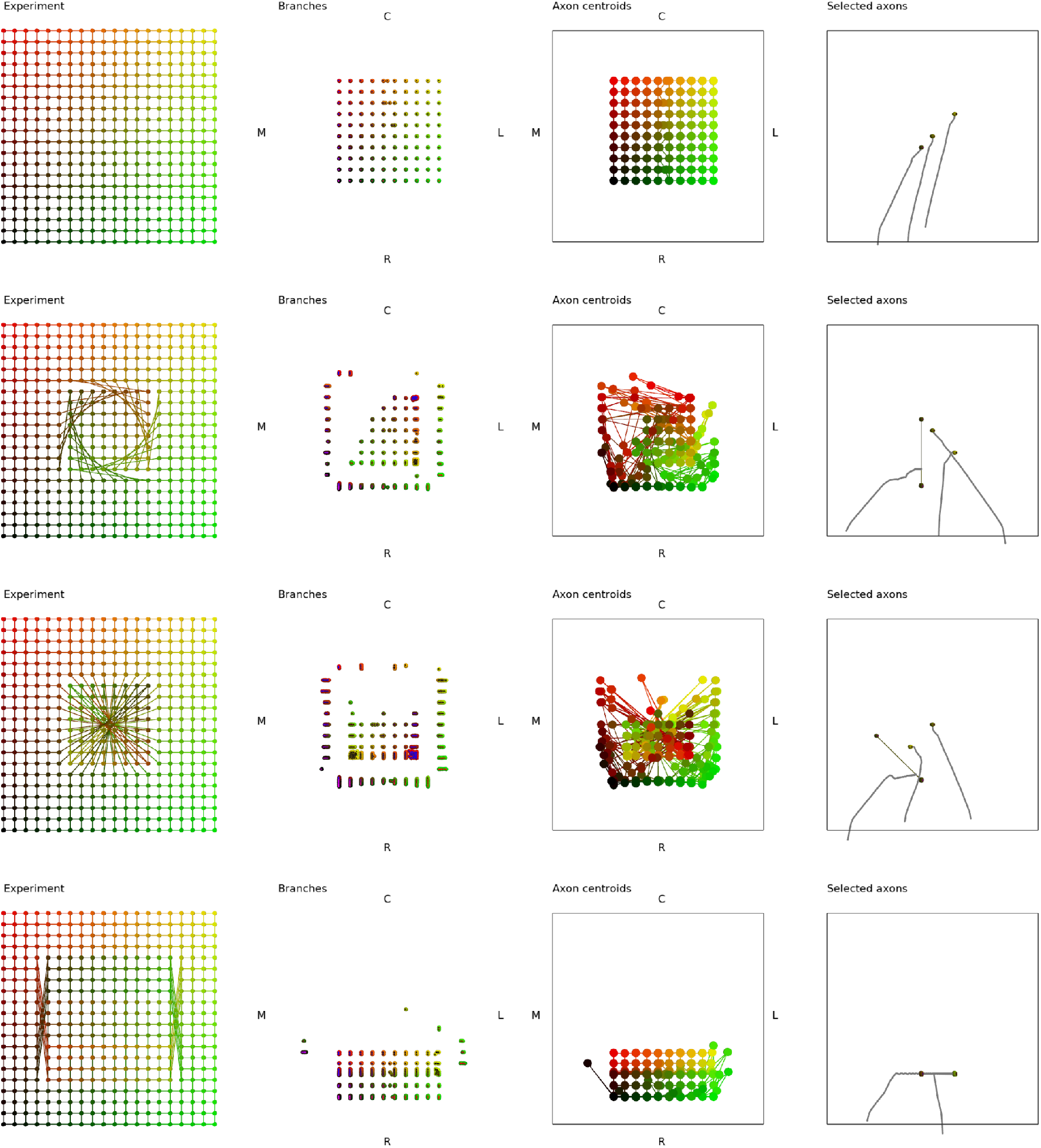
Surgical manipulations in the chemoaffinity-only model. Row A: wildtype. Row B: 90 degree rotation. Row C: 180 degree rotation. Row D: Graft swap

### Grating simulations

I’d like to do this if it’s not too much work!

## 4. Discussion

The model described above assumes that branches express graded levels of receptors which interact with ligands expressed as gradients on the tectum, the tectal border and on the surface of other branches, and that only these interactions guide their movement.

